# Detection of Antibiotic-resistant *Escherichia coli* in the Euphrates River, Iraq

**DOI:** 10.1101/2025.08.30.673253

**Authors:** Zainab Altameemi, Adel Talib, Saad Fakhry

## Abstract

Water plays a crucial role in the spread of antibiotic-resistant bacteria (ARB) among humans, animals, and the environment. The emergence of antibiotic resistance is a global health and environmental problem. This study aimed to identify the temporal and spatial patterns and hotspots of antibiotic-resistant *Escherichia coli* (AREs) in the Euphrates River in central Iraq. Integrating *Escherichia coli* (*E. coli*) testing with key environmental parameters allowed for linking resistant bacteria to pollution sources. Water samples were collected from four selected points along the river during the dry and wet seasons of 2023 and 2024. Fecal samples (non-environmental samples) were also collected from areas surrounding the river to study their impact on it. Resistance patterns of *E. coli* and multidrug resistance to 12 antibiotics of 11 rows were tested for 80 river water isolates and 39 non-environmental isolates by using methodological methods. *E. coli* showed the highest resistance to ampicillin (97.5%), followed by cefotaxime (96.25%). Importantly, multidrug resistance was recorded in 93.75% and 97.44%, of river and non-environmental isolates, respectively. The recorded values for the Antibiotic Resistance Index exceeded the threshold. Environmental parameters showed varying degrees of association with antibiotic resistance. There are few reports about ARB in Iraqi rivers due to the focus on clinical samples. Nevertheless, a full understanding of the origin, spread, and persistence of antibiotic resistance requires an integrative strategy that considers all ecosystems that may contribute to its spread. Our findings demonstrate the urgent need to address fecal contamination and the spread of ARB in Iraqi rivers.

## 1. Introduction

Water quality is assessed by chemical, physical, and biological parameters [1], [2]. Physicochemical parameters including temperature, electrical conductivity, total dissolved solids, turbidity, dissolved oxygen, and pH levels are often used indicators of water quality. A physicochemical and bacteriological examination is necessary to adequately examine the many techniques and processes involved in contaminating freshwater bodies [3]. Water is seldom found in its purest form in nature since it is frequently contaminated by human waste, pollutants, and the introduction of harmful organisms [4]. Water quality is a worldwide concern, with unimproved contaminated water sources improper sanitation practices lead to human diseases [5]. Rivers as one of essential natural water resources that have significant impacts on the public health, economy, and food production [6]. It is well known that a wide range of microbiological pathogens, such as bacteria, viruses, parasites, and chemical pollutants, can be found in contaminated water [7]. Antibiotic residues pollute soil and water sources, which may eventually pose a hazard to human health and have negative environmental repercussions, including the emergence of resistant microbes [8]. A study conducted for 258 rivers in the world showed that concentrations of active pharmaceutical substances exceed safe limits in more than a quarter of them [9]. Antibiotics’ physicochemical qualities, partition features, and environmental behavior all influence the presence and destiny of antibiotic residues in aquatic environments [10].

As one of the top ten dangers to world health, ARB is highlighted by the World Health Organization (WHO), which supports the “One Health” concept, which calls for immediate coordinated action to address the health of human, animal, and the environment. In particular, the WHO warns that we might be entering a post-antibiotic era in which common medical procedures that depend on preventative antibiotic treatment may no longer be feasible and basic human infections could once again become fatal [11]. Recent research has recognized the natural environment as a potentially significant source of ARB infections to humans as well as animals [12], which enter water from various routes, such as hospital wastewater, agricultural wastewater, and aquaculture wastewater [13]. The ARBs have the potential to re-enter the environment and infiltrate individuals who utilize the water source for drinking, recreation, swimming, or water sports, as well as the food chains. In this sense, ARB is disseminated and stored in the aquatic environment [14]. Rivers play an important role in the transfer of ARB between different environments. They may be a potential pathway for the transfer of ARB between the environment, animals, and humans [15]. However, national and international authorities responsible for monitoring water quality have not included ARB in their contaminant monitoring lists. For instance, they are not listed as contaminants in the US Water Quality Criteria list or in the European Water Framework Directive of pollutants [16].

Investigations on the prevalence and forms of ARB in freshwater are scarcer than in hospitals and therapeutic settings, and the environmental community has overlook this issue [17]. The intricacy of this potentially worldwide problem is further compounded by the possibility that bacteria will develop ARB to several antibiotic classes, which would increase the prevalence of multidrug resistance (MDR) in populations of bacteria, especially potential pathogens [18]. ARB pathogens are directly responsible for nearly a million deaths per year [19]. Since 10 million fatalities a year are predicted to be linked to ARB by 2050, the overuse and abuse of antibiotics in human and animal medicine, as well as the levels of residual antibiotics discharged into the environment via wastewaters has made ARB a serious issue [20], [21]. At the same time, six billion people are expected to suffer from severe water shortages by 2050, according to UN Water Development estimate [22].

*E. coli* is a member of the environment’s bacterial, as well as commensal bacteria in the gut of animals and humans, making it a reliable indicator of fecal contamination. It also includes highly pathogenic strains that cause serious disease outbreaks [23]. It serves as the main source of genes that cause antibiotic resistance and are readily transmitted to pathogenic bacteria [24], [25]. Waterborne *E. coli* that is resistant to antibiotics is a global health concern, particularly in developed nations. AREs from diverse habitats must be continuously monitored in order to preserve public health [26].

To our knowledge, the diversity and prevalence of antibiotic resistance factors of *E. coli* isolates in Iraqi rivers are still rare. Therefore, the aim of this study was to investigate and monitor AREs in the water of the Euphrates River and animal feces from agricultural lands surrounding the river in central Iraq, in addition detecting environmental factors associated with ARB pollution in rivers. This study documents the potential threat to human, animal, and environmental health in the Euphrates River posed by ARB and multidrug resistance, paving the way for future monitoring and remediation efforts.

## 2. Results and Discussion

### 2.1. Physicochemical Parameters

The results were presented in another study submitted for publication, although some parameters exceeded standard guidelines.

### 2.2. Identification *E. coli*

In this study, AREs in Euphrates River water were characterized to understand their potential impact on public health. A total of 112 colonies of potential *E. coli* were isolated from the river water and subjected to screening. Biochemical and polymerase chain reaction (PCR) analysis using specific primers targeting the *uidA* housekeeping gene (Figure 1) confirmed that 71.4% (80/112) of the isolates were *E. coli*. 45 isolates returned to the wet season and 35 to the dry season. As for the sites, 16 isolates were distributed to S1, 20 to S2, 17 to S3, and 27 to S4. Returning to the non-environmental samples after conducting the same previous tests, 78% (39/50) isolates were confirmed to contain *E. coli*.

**Figure 1.**
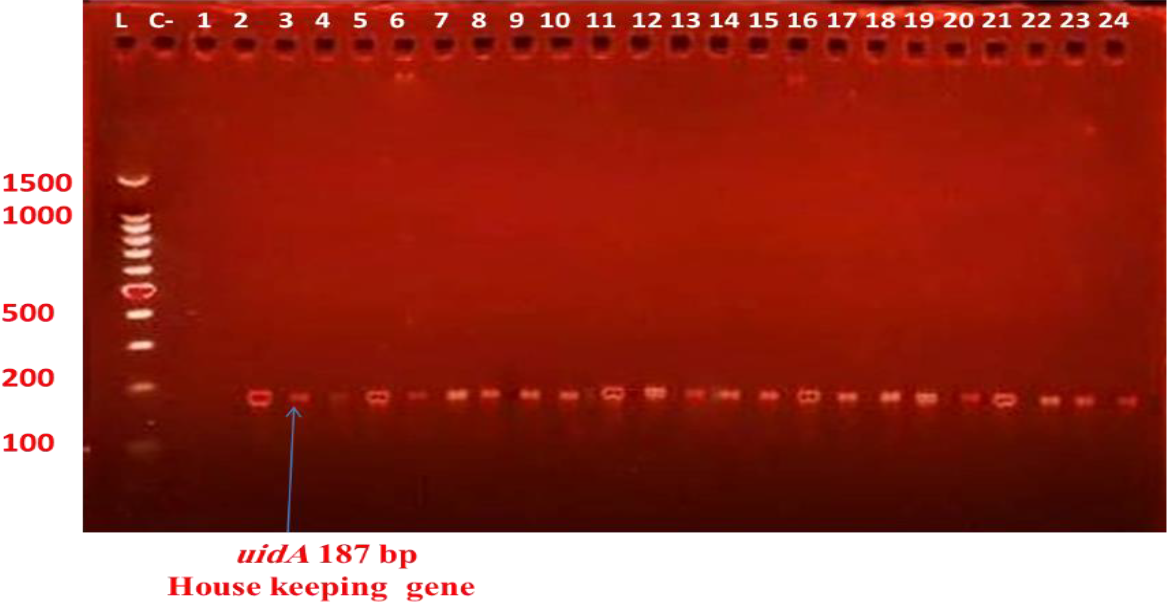
Products of PCR Lane L; DNA Ladder, Lane C-; Negative control, Lanes 1-24; positive *E.coli*.

### 2.3. Antibiotic Resistance Phenotypes of *E. coli*

Twelve antibiotics were used to test the antibiotic resistance of the *E. coli* isolates from Euphrates River (n=80). Resistance-designated isolates exhibit phenotypic resistance to one or two antibiotic classes, while multi-resistant isolates exhibit resistance to three or more. High levels of resistance were observed isolates subjected to antibiotic susceptibility testing. The highest resistance rate was recorded for AMP 97.5% (78/80), followed by CTX 96.25% (77/80), while the least resistance was to MRP 21.25% (17/80) (Figure 2, 5). Our results showed that *E. coli* remained sensitive to MRP 68.75% (55/80), followed by CTR 48.75% (39/80), while no sensitivity was recorded towards either AMP or NA 0% (Figure 3, 6). The highest value of intermediate resistance was recorded to C 38.75% (31/80), followed by NA 36.25% (29/80), while no intermediate resistance was recorded towards AZM 0% (Figure 2, 5).

**Figure 2.**
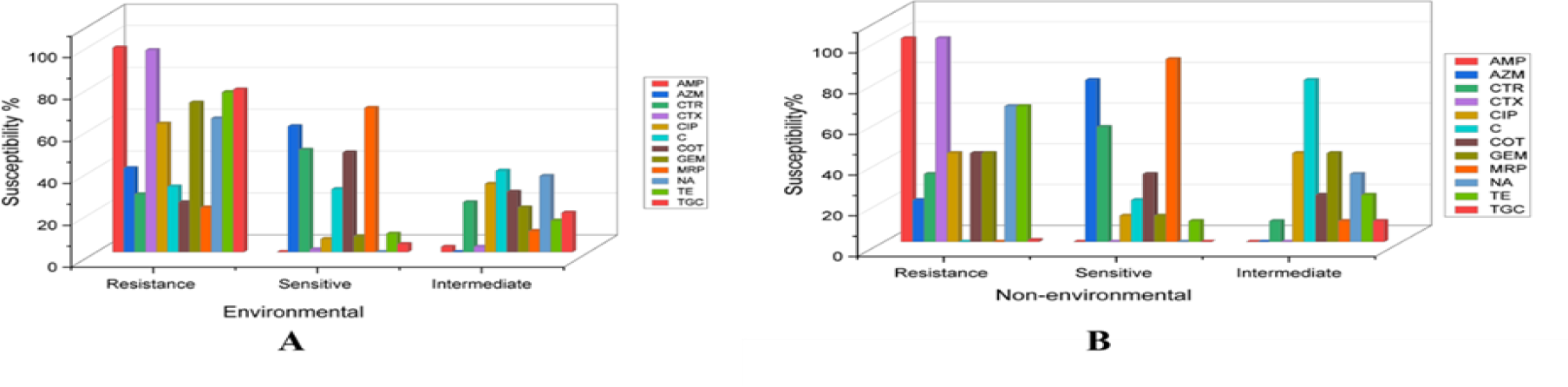
Antibiotics sensitivity pattern of *E. coli* isolates from (A) the Euphrates Rivers (B) the non-environmental.

**Figure 3.**
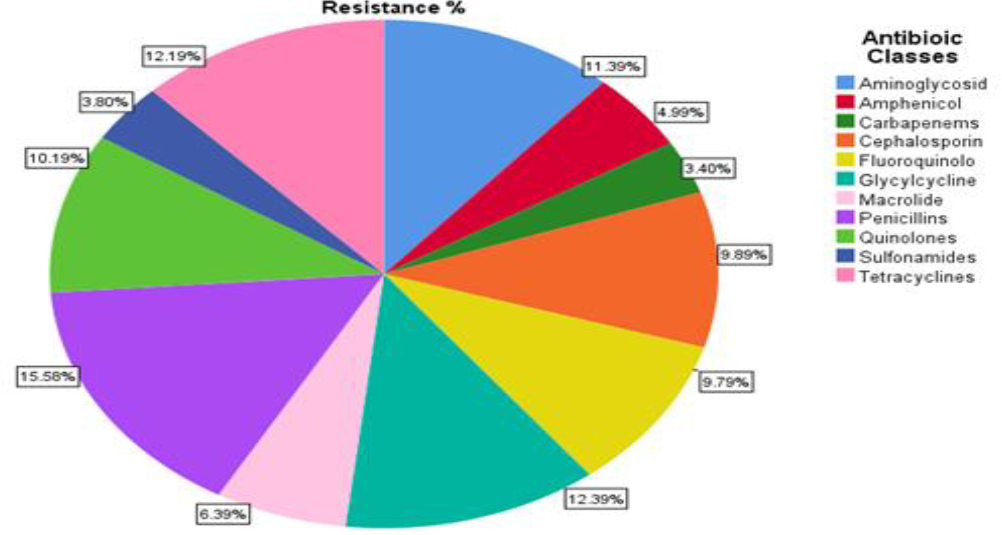
Antibiotic resistance percentage according to antibiotic classes of E. *coli* isolates from the Euphrates River.

According to antibiotic classes, the highest resistance rate was recorded for penicillins 97.5 %, followed by glycylcycline 77.5%, and tetracyclines 76.25% (Figure 3). All isolates were resistant to at least two antibiotic class, while two *E. coli* strains showed resistance to all antibiotic classes. Differences in antibiotic resistance according to study sites and seasons are shown in the (Figure 4).

**Figure 4.**
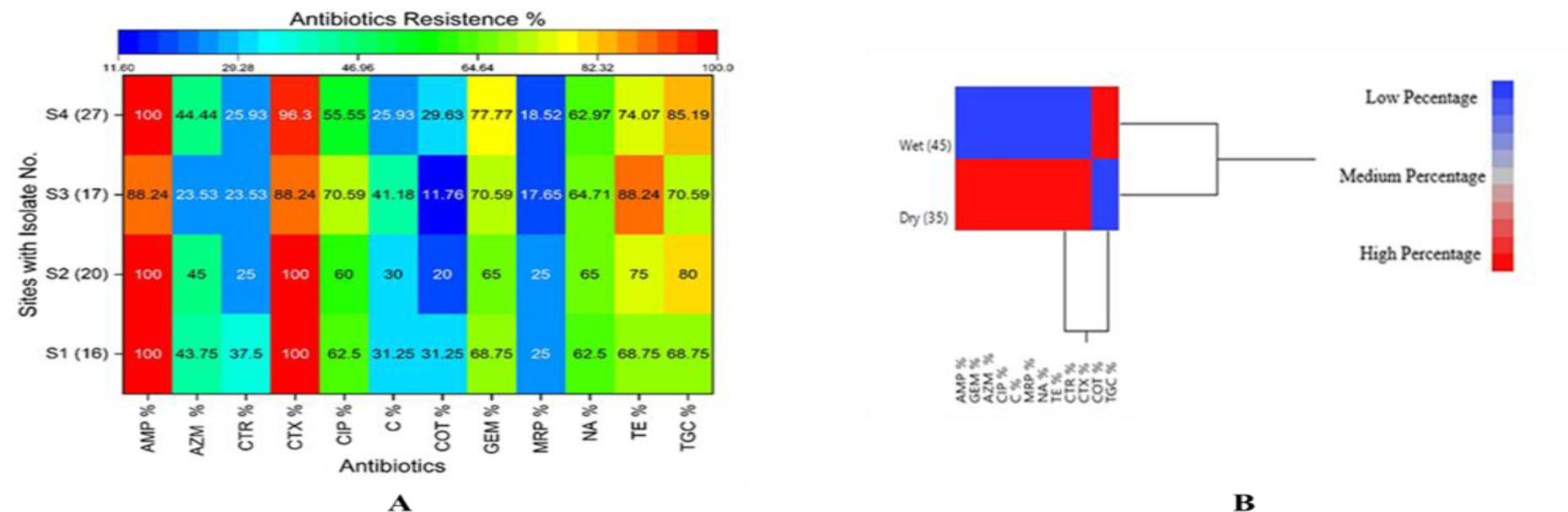
Percentage of antibiotic resistance according to (A) study sites and (B) study seasons.

**Figure 5.**
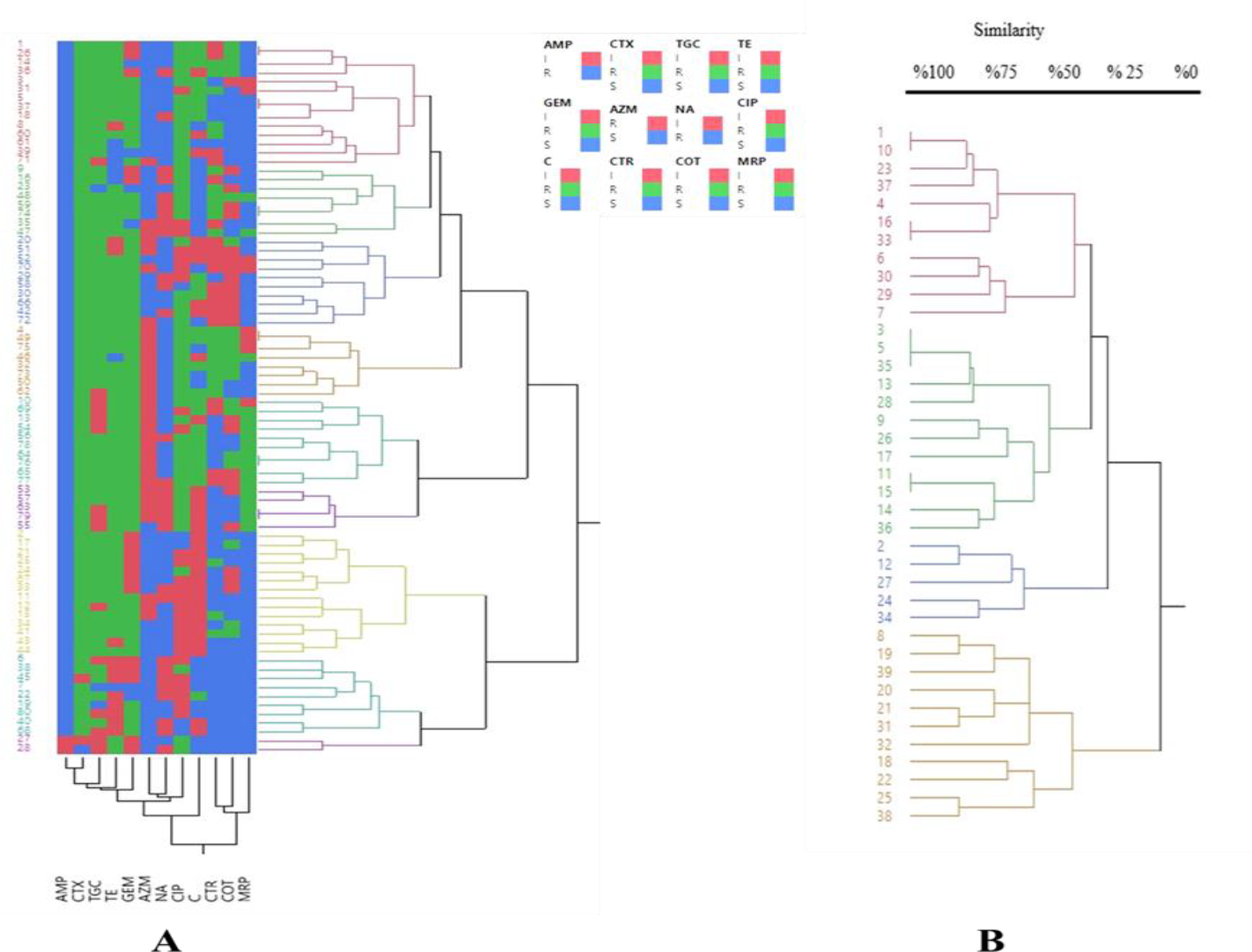
A hierarchical cluster heat map of antibiotic resistance patterns of *E. coli* isolates from (A) the Euphrates River and (B) non-environmental isolates.

In non-environmental isolates, the highest resistance was towards both AMP and CTX 100%, followed by TGC 89.74% (35/39), while the lowest resistance was towards AZM 20.51% (8/39). No resistance was recorded towards MRP and C. The highest sensitivity was recorded towards MRP 89.74% (35/39), and the highest intermediate resistance was towards AZM 76.92% (30/39) (Figure 3, 6). Analysis of variance (ANOVA) test did not show significant differences in antibiotic resistance between river isolates and non-environmental isolates except for (AMP-C, AMP-MRP, C-CTX, CTX-MRP) *p* < 0.05.

**Figure 6.**
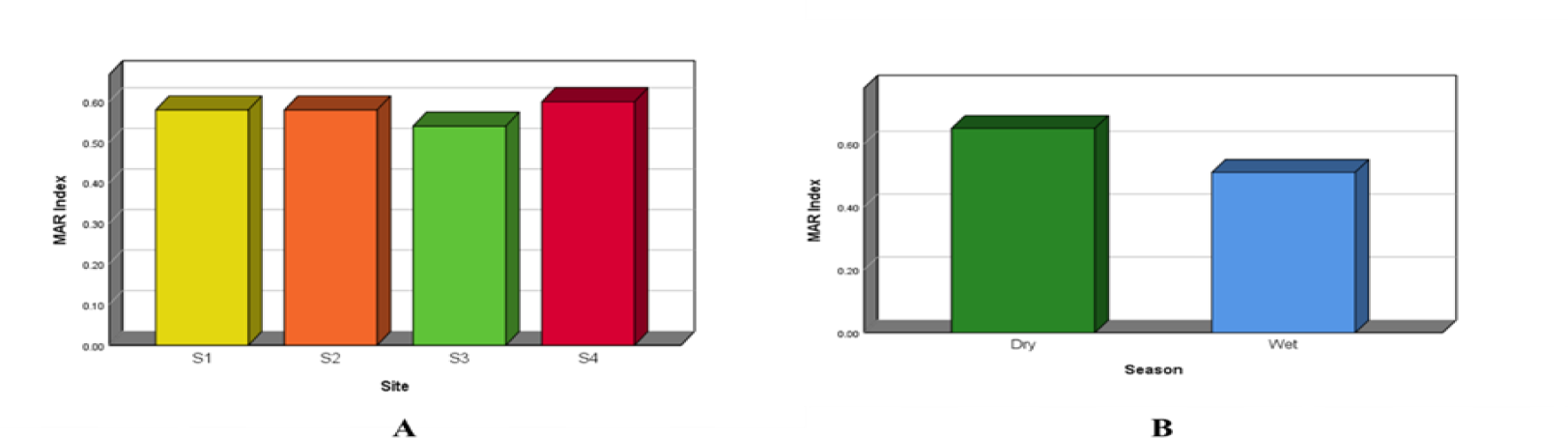
Spatial (A) and Temporal (B) MAR values of *E. coli* isolates (n=80) from the Euphrates River.

**Figure 7.**
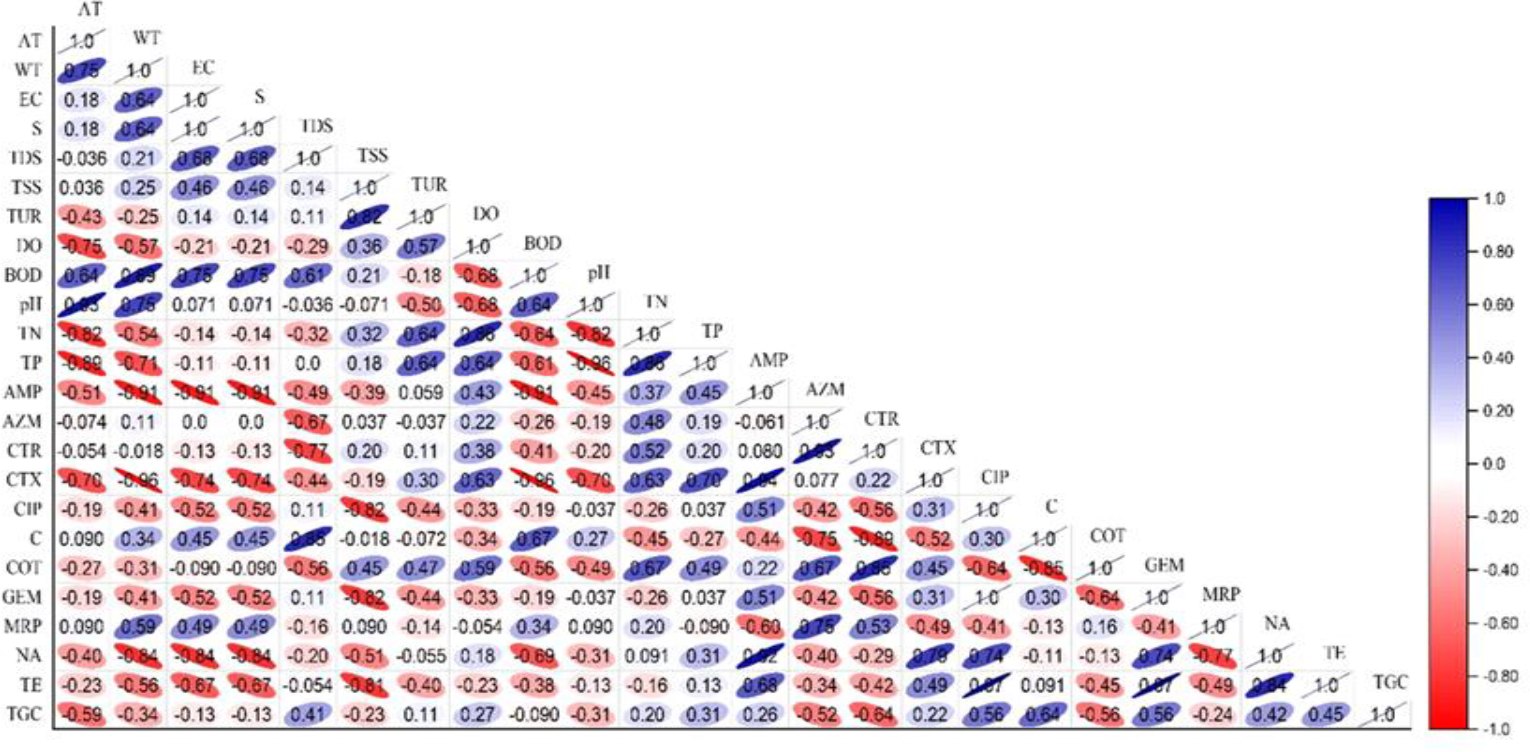
Association matrix between AREs and environmental parameters.

#### 2.3.1. Multidrug-resistant of *E. coli* (MDR)

The rate of multidrug-resistant MDR were identified as 93.75% (70/80) in different patterns (Table 1). More importantly, the results of this study showed that 53.75% (43/80) of the isolates in this study were resistant to more than six classes of antibiotics. Among the multidrug-resistant isolates, 21.25% were classified as carbapenem-resistant. For non-environmental isolates, MDR was recorded at 97.44%, some patterns similar to those of river isolates. The most similar pattern between river and non-environmental isolates was (AMP, CTR, CTX, COT, GEM, NA, TE, TGC). The highest resistance to antibiotic classes was recorded in human isolates.

**Table 1.**
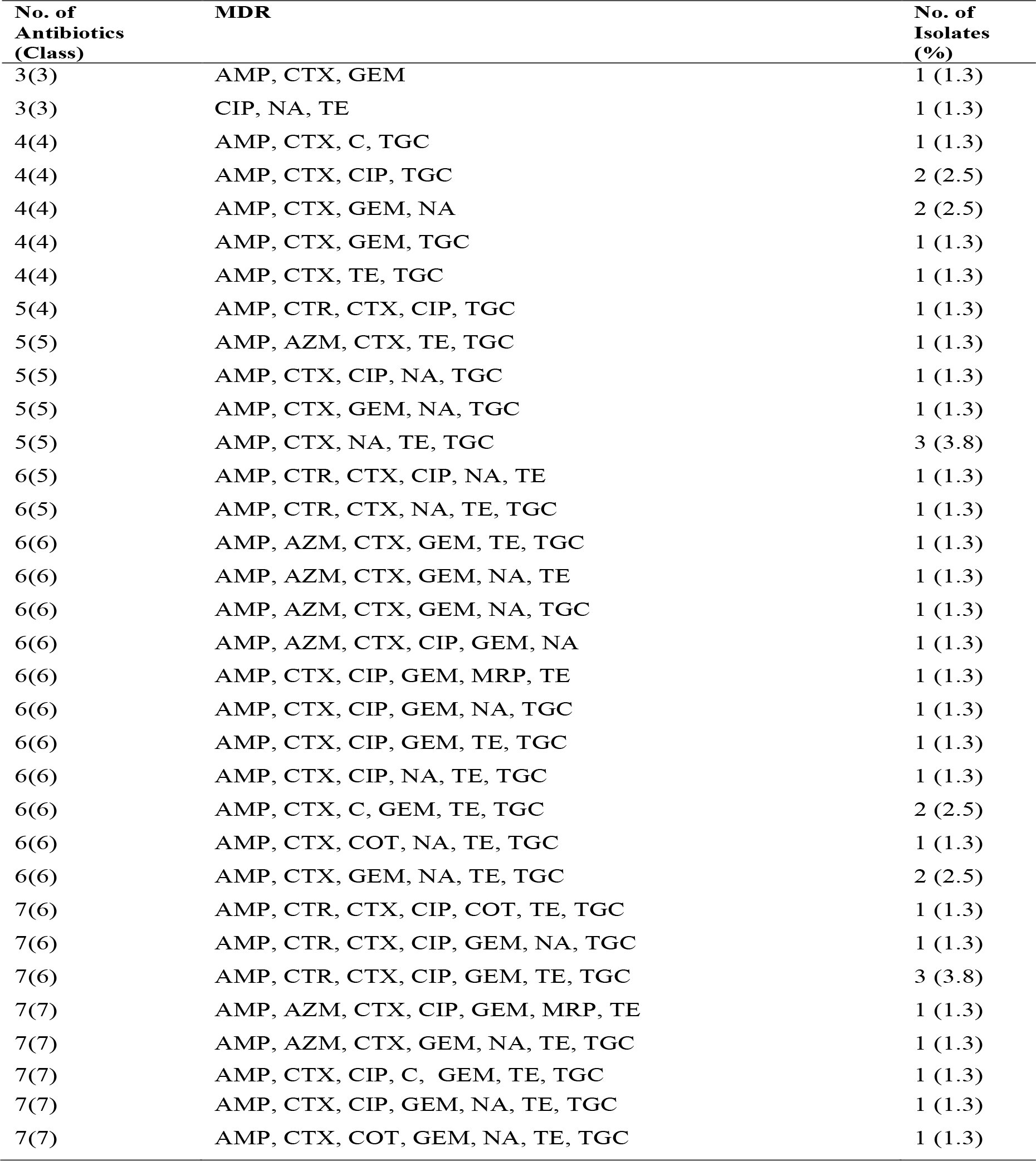
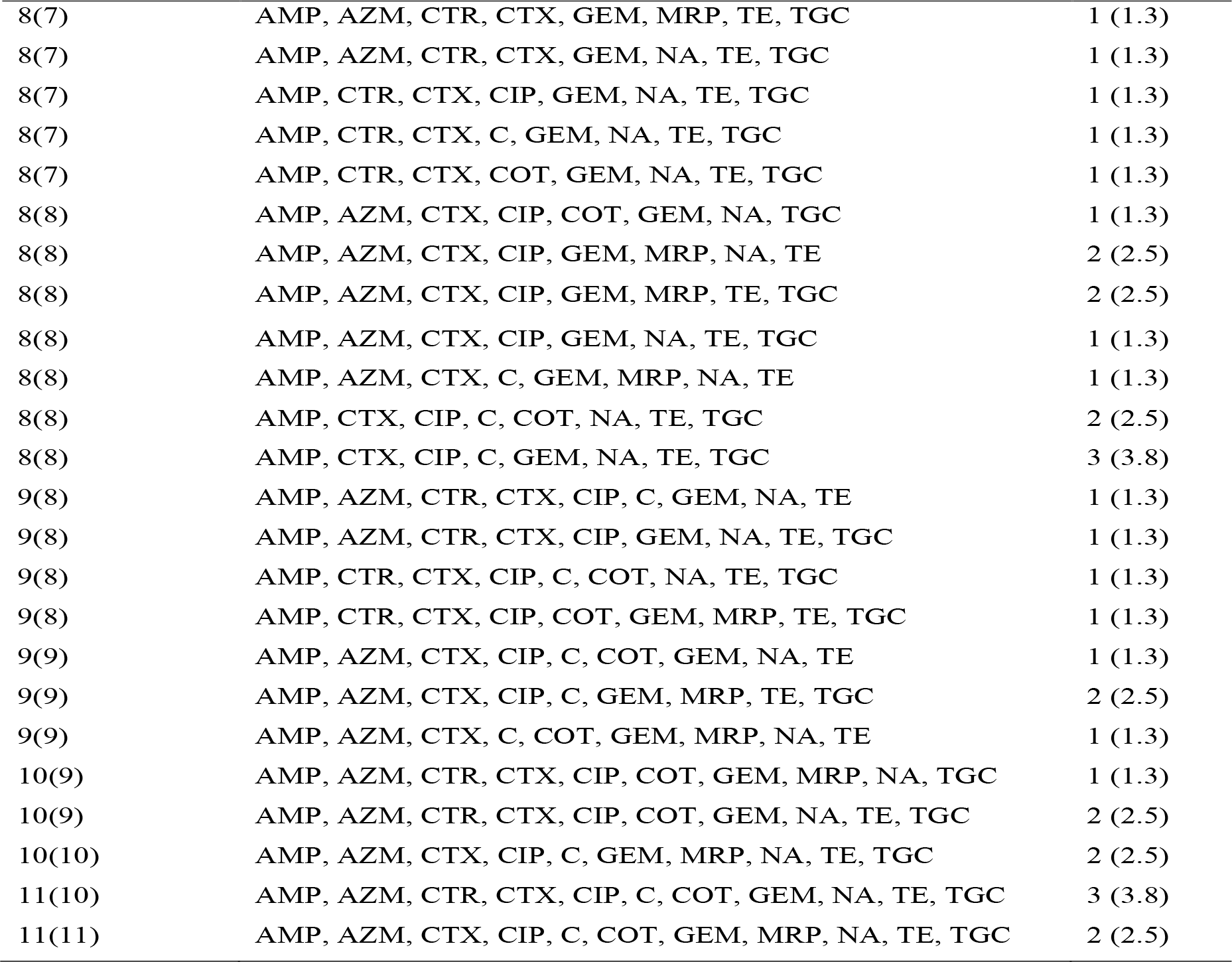
Multidrug resistance (MDR) pattern of *E. coli* isolates (n= 64) from the Euphrates River.

| **No. of Antibiotics (Class)** | **MDR** | **No. of Isolates (%)** |
| --- | --- | --- |
| 3(3) | AMP, CTX, GEM | 1 (1.3) |
| 3(3) | CIP, NA, TE | 1 (1.3) |
| 4(4) | AMP, CTX, C, TGC | 1 (1.3) |
| 4(4) | AMP, CTX, CIP, TGC | 2 (2.5) |
| 4(4) | AMP, CTX, GEM, NA | 2 (2.5) |
| 4(4) | AMP, CTX, GEM, TGC | 1 (1.3) |
| 4(4) | AMP, CTX, TE, TGC | 1 (1.3) |
| 5(4) | AMP, CTR, CTX, CIP, TGC | 1 (1.3) |
| 5(5) | AMP, AZM, CTX, TE, TGC | 1 (1.3) |
| 5(5) | AMP, CTX, CIP, NA, TGC | 1 (1.3) |
| 5(5) | AMP, CTX, GEM, NA, TGC | 1 (1.3) |
| 5(5) | AMP, CTX, NA, TE, TGC | 3 (3.8) |
| 6(5) | AMP, CTR, CTX, CIP, NA, TE | 1 (1.3) |
| 6(5) | AMP, CTR, CTX, NA, TE, TGC | 1 (1.3) |
| 6(6) | AMP, AZM, CTX, GEM, TE, TGC | 1 (1.3) |
| 6(6) | AMP, AZM, CTX, GEM, NA, TE | 1 (1.3) |
| 6(6) | AMP, AZM, CTX, GEM, NA, TGC | 1 (1.3) |
| 6(6) | AMP, AZM, CTX, CIP, GEM, NA | 1 (1.3) |
| 6(6) | AMP, CTX, CIP, GEM, MRP, TE | 1 (1.3) |
| 6(6) | AMP, CTX, CIP, GEM, NA, TGC | 1 (1.3) |
| 6(6) | AMP, CTX, CIP, GEM, TE, TGC | 1 (1.3) |
| 6(6) | AMP, CTX, CIP, NA, TE, TGC | 1 (1.3) |
| 6(6) | AMP, CTX, C, GEM, TE, TGC | 2 (2.5) |
| 6(6) | AMP, CTX, COT, NA, TE, TGC | 1 (1.3) |
| 6(6) | AMP, CTX, GEM, NA, TE, TGC | 2 (2.5) |
| 7(6) | AMP, CTR, CTX, CIP, COT, TE, TGC | 1 (1.3) |
| 7(6) | AMP, CTR, CTX, CIP, GEM, NA, TGC | 1 (1.3) |
| 7(6) | AMP, CTR, CTX, CIP, GEM, TE, TGC | 3 (3.8) |
| 7(7) | AMP, AZM, CTX, CIP, GEM, MRP, TE | 1 (1.3) |
| 7(7) | AMP, AZM, CTX, GEM, NA, TE, TGC | 1 (1.3) |
| 7(7) | AMP, CTX, CIP, C, GEM, TE, TGC | 1 (1.3) |
| 7(7) | AMP, CTX, CIP, GEM, NA, TE, TGC | 1 (1.3) |
| 7(7) | AMP, CTX, COT, GEM, NA, TE, TGC | 1 (1.3) |
| 8(7) | AMP, AZM, CTR, CTX, GEM, MRP, TE, TGC | 1 (1.3) |
| 8(7) | AMP, AZM, CTR, CTX, GEM, NA, TE, TGC | 1 (1.3) |
| 8(7) | AMP, CTR, CTX, CIP, GEM, NA, TE, TGC | 1 (1.3) |
| 8(7) | AMP, CTR, CTX, C, GEM, NA, TE, TGC | 1 (1.3) |
| 8(7) | AMP, CTR, CTX, COT, GEM, NA, TE, TGC | 1 (1.3) |
| 8(8) | AMP, AZM, CTX, CIP, COT, GEM, NA, TGC | 1 (1.3) |
| 8(8) | AMP, AZM, CTX, CIP, GEM, MRP, NA, TE | 2 (2.5) |
| 8(8) | AMP, AZM, CTX, CIP, GEM, MRP, TE, TGC | 2 (2.5) |
| 8(8) | AMP, AZM, CTX, CIP, GEM, NA, TE, TGC | 1 (1.3) |
| 8(8) | AMP, AZM, CTX, C, GEM, MRP, NA, TE | 1 (1.3) |
| 8(8) | AMP, CTX, CIP, C, COT, NA, TE, TGC | 2 (2.5) |
| 8(8) | AMP, CTX, CIP, C, GEM, NA, TE, TGC | 3 (3.8) |
| 9(8) | AMP, AZM, CTR, CTX, CIP, C, GEM, NA, TE | 1 (1.3) |
| 9(8) | AMP, AZM, CTR, CTX, CIP, GEM, NA, TE, TGC | 1 (1.3) |
| 9(8) | AMP, CTR, CTX, CIP, C, COT, NA, TE, TGC | 1 (1.3) |
| 9(8) | AMP, CTR, CTX, CIP, COT, GEM, MRP, TE, TGC | 1 (1.3) |
| 9(9) | AMP, AZM, CTX, CIP, C, COT, GEM, NA, TE | 1 (1.3) |
| 9(9) | AMP, AZM, CTX, CIP, C, GEM, MRP, TE, TGC | 2 (2.5) |
| 9(9) | AMP, AZM, CTX, C, COT, GEM, MRP, NA, TE | 1 (1.3) |
| 10(9) | AMP, AZM, CTR, CTX, CIP, COT, GEM, MRP, NA, TGC | 1 (1.3) |
| 10(9) | AMP, AZM, CTR, CTX, CIP, COT, GEM, NA, TE, TGC | 2 (2.5) |
| 10(10) | AMP, AZM, CTX, CIP, C, GEM, MRP, NA, TE, TGC | 2 (2.5) |
| 11(10) | AMP, AZM, CTR, CTX, CIP, C, COT, GEM, NA, TE, TGC | 3 (3.8) |
| 11(11) | AMP, AZM, CTX, CIP, C, COT, GEM, MRP, NA, TE, TGC | 2 (2.5) |

#### 2.3.2. Multiple antibiotic resistance (MAR) index

Our results showed that the MAR values for the study sites and seasons all exceeded the 0.2 threshold (S4 > S1 = S2 > S3), (dry > wet) (Figure 6), indicating a high incidence of contamination of river water with AREs. Non-environmental isolates were similar to river isolates in that the index values were high, reaching 0.51.

### 2.4. Correlation between AREs and Environmental Parameters

AREs showed varying degrees of association with environmental parameters. In general, a negative relationship was found, except for TN and TP, which were positively associated (Figure 8).

**Figure 8.**
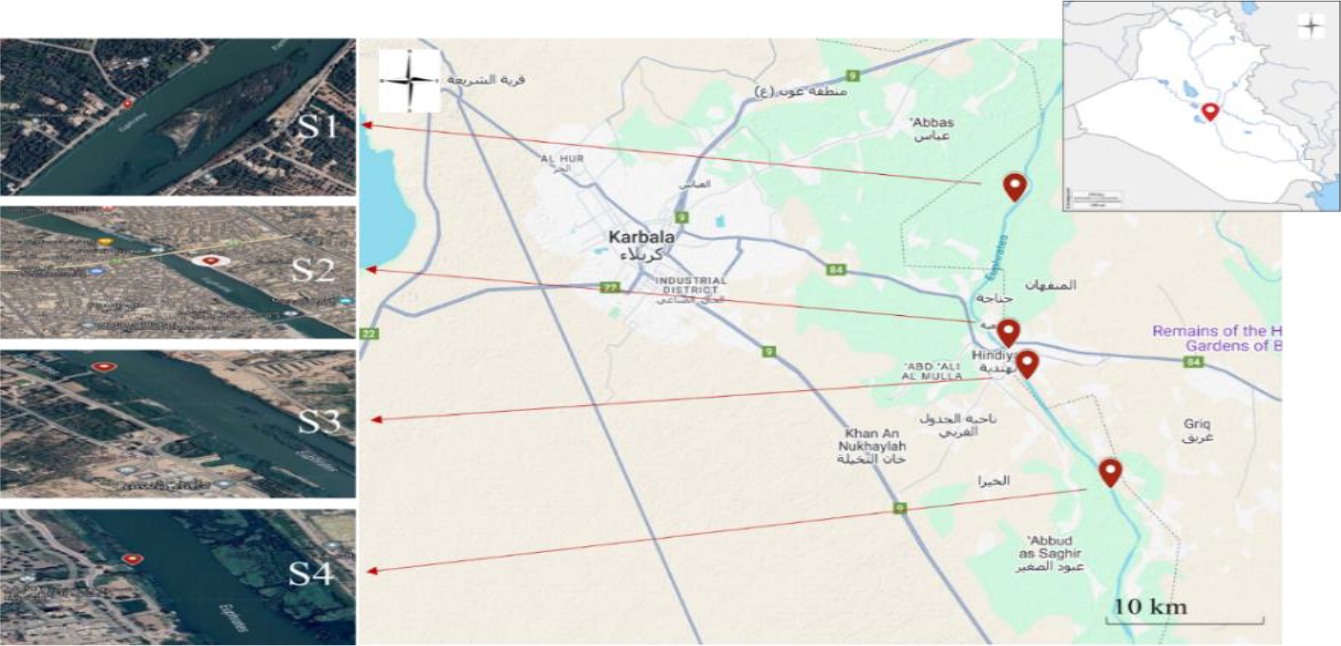
An interactive map that allows clicking on dots to show the sites of study in Euphrates River (Supplement).

The global prevalence of ARB in the environment poses a growing health threat. Rivers are of particular concern due to their high vulnerability to wastewater discharge, yet they represent a vital lifeline that meets diverse human needs [27]. AREs in aquatic habitats is thought to be a reliable indicator of antibiotic pollution from sewage or animal waste contamination [26]. In order to identify environmental pollution and “hotspots” of ARB, it is crucial to determine the spatial distribution of significant ARB [28]. In recent decades, cities along the Euphrates River have been suffering high levels of pollution, especially downstream of the river [29], also a decrease in the level of surface runoff due to the construction of dams by upstream countries [30]. Intensive agricultural activities, untreated wastewater with an increase in the number of coliform bacteria and nutrients, increased evaporation rates, strong climate fluctuations, poor drainage, and the accumulation of salts and sediments are factors that led to a decrease in the quality of the river’s water and an increase in the level of pollution in it [31].

The most common and clinically significant antibiotic-resistant Gram-negative bacteria include *E. coli*, which is classified as pathogen on the WHO list of potentially fatal diseases [32]. Antibiotic resistance makes treating infectious diseases more difficult, raises financial expenses, lengthens the course of therapy, raises financial expenses, and raises fatality rates. Therefore, resistant infections pose a serious threat to human, animal, and environmental health [33].

The highest levels of resistance in this study were recorded against ampicillin, which is a semisynthetic antibiotic belonging to the aminopenicillin group. It is used as a valuable agent in the treatment of certain bacterial infections, such as *E. coli*, where resistance is the most common [34]. The results of this study are consistent with those of the World Health Organization in Europe, which demonstrated clinical *E. coli* resistance to aminopenicillin [35], signifying a concurrent rise in the resistance rates of both clinical and environmental *E. coli*. Many previous study has also recorded high levels of *E. coli* resistance to ampicillin [36], such as 97% in the surface water in the Irish [37]. Ampicillin was the antibiotic most resistant to *E. coli* in the following studies, but at less than what was recorded in our results: in two rivers in Taiwan [26], surface water in Belgium [38], the Danube River in Austria [39], the Erkyna river and the Boeotian river in Greece [40]. Our results found isolates with a high percentage of resistance to last-line antibiotics, such as tigecycline 77.5%, which poses a major threat due to the lack of treatment for pathogens resistant to the latest class of antibiotics [41], [42]. Recent studies have recorded the presence of *E. coli* resistant to tetracycline in surface water and wastewater, but at a lower rate than recorded in our results [43], [41], [39]. Resistance to tetracycline and has been observed. A possible explanation for this is the common use of tetracycline in veterinary medicine (Kumar et al., 2019). Our results showed that resistance to ciprofloxacin reached 61.25%, which is a widely used drug found in a variety of environmental matrices, including groundwater, surface waterways, and soil. The high concentration of ciprofloxacin in water (estimated at 50–80%) is the cause of the high resistance to the antibiotic. This is due to the fact that ciprofloxacin is widely used in aquaculture to treat and prevent bacterial infections in aquatic animals. Furthermore, this chemical may enter the environment by a number of routes, including wastewater discharges, agricultural runoff, and landfill leachate [44], [45], [45], [46]. The persistent nature of ciprofloxacin in wastewater and its ability to persist in the environment for long periods of time may cause environmental microorganisms to become resistant to antibiotics [47], [48]. A study of the upper Euphrates River in Anbar Province, Iraq, showed that levels of *E. coli* resistance to ciprofloxacin, ceftriaxone, and meropenem were higher than recorded in our results [49]. While resistance against cefotaxime, ciprofloxacin, and Gentamicin in a study of the Tigris River in Iraq was less than what was recorded in this study [50]. Azithromycin belongs to the macrolide class of antibiotics, which are described by their large lactone ring structure, making them effective in treating intestinal bacterial infections, including *E. coli* [51]. The results of this study showed the prevalence of resistance towards the classes of antibiotics penicillins, tetracyclines, and glycylcycline, respectively. Our results are consistent with other studies that have reported resistance of *E. coli* to older and widely used antibiotics in aquatic environments [52], [53], [54], [39]. Carbapenem class resistance was low 21.25% (Figure 4). This should be regarded as a warning indication, though, as carbapenems are antibiotics used as a last resort to treat complex, potentially fatal infections in humans [6].

Multidrug-resistant bacteria are a concern due to the increasing ARB, its potential impact on patient safety, and its effects on surveillance systems [55], [56]. If inadequately managed, the concerning incidence of multidrug resistant *E. coli* could result in dire public health implications [57]. This study observed high levels of multidrug resistance in *E. coli* with varying patterns, which is consistent with a local study of the Tigris River, Baghdad, Iraq [58], and several global studies, including one on the Kshipra River in India [15], in Meia Ponte River in Brazil [17], in the Douro River in Portugal [59], in the Themi River in Tanzania [60]. Our results for non-environmental isolates are consistent with a study showing high antibiotic resistance in *E. coli* isolates from wild and domestic animals, and they are considered potential sources of multidrug resistance. High levels of AREs have also been linked to domestic animals [61].

The high MAR index at site 4 may be attributed to the increased number of isolates at the site and its rural and agricultural nature. Studies have shown that livestock and poultry farms have high levels of ARB and are reservoirs of ARB [62], [28], [63]. Animal dung that washes into the water is also a hotspot for these bacteria. In Site 2, the high index is due to wastewater treatment plant discharges, a proven source of ARB [64]. The rise in the MAR index during the dry season may be due to the increase in human recreational activities during this season [65]. The environmental MAR values detected in this study was comparable to those of the Vembanad Lake study in India [66], but higher than the Larut River in Malaysia [67].

Antibiotic resistance rates were inversely related to pH, and positively related to TN and TP, which is consistent with a study conducted on the Yellow River in China [68]. Various factors can influence the concentrations of antibiotics and their resistance genes in rivers, including factors associated with climate change (such as temperature, water scarcity, dissolved organic carbon, and water dilution) and the amount of antibiotics and biocides in wastewater from urban wastewater treatment plants and hospitals [16]. The temperature, alkaline, and pH conditions might impact the hydrolysis and degradation of antibiotics, whereas total dissolved solids may play a role in the selection procedure for antibiotic resistance [15].

Comparing the Euphrates River isolates with the non-environmental isolates, both showed high levels of antibiotic resistance, with no significant differences recorded in most of them. Hierarchical clustering also showed convergence between them. The results of the Pearson test showed a strong positive correlation (r= 0.883, *p* < 0.001) between the river isolates and the non-environmental isolates. Regarding the patterns of multi-antibiotic resistance, both isolates showed convergence, with a higher rate of multi-resistance recorded in the non-environmental isolates. This indicate the presence of a pathway for the transmission of AREs from the feces of animals and humans, which are its reservoirs, to the environment [69]. Human activities, such as agriculture, increase the load of environmental bacteria resistant to antibiotics. Reports have indicated the presence of diverse resistance genes in agricultural environments and increased levels of antibiotic resistance genes in soil after the application of organic manure [70], which is washed into surface and groundwater from lands adjacent to the river [71].

## 3. Materials and Methods

### 3.1. Study Site Description

The Euphrates is the longest river in western Asia, with a total length of 2940 km, and 1213 km of distance in Iraq about 40% [72]. The catchment area 440,000 km^2^ covers the Euphrates basin, with 47% of that area located in Iraq [73]. The Euphrates River constitutes the basis of life in Iraq, as one-third of Iraq’s population lives on its banks, and it penetrates eight Iraqi governorates. Agriculture also depends entirely on irrigation due to the fluctuation and scarcity of rain [74]. We selected four diverse sites along the Euphrates River in Karbala Province, Iraq, which include agricultural, livestock, and urban activities, to collect study samples. The selection of the study locations followed the recommendations of [75]. The first site (S1) is located after the Hindiyah Dam (32°37’29.9”N, 44°13’48.2”E), while the second site (S2) is located in the centre of the Hindiyah city with a high population density, near the water treatment plant discharge and recreational facilities on its banks (32°32’41.0”N, 44°13’35.2”E). Followed by the third site (S3) after the city of Hindiyah (32°31’39.3”N, 44°14’12.8”E). Finally, the fourth site (S4) in agricultural lands (32°28’04.7”N, 44°17’02.2”E) (Figure 8).

### 3.2. Water and *E. coli* Sampling

Samples were collected seasonally from November 2023 to September 2024. From four chosen locations, duplicate water samples from the river have been collected in sterile containers. A total of 1.5 L of water has been collected from every sampling location for physicochemical assessments, 500 mL to isolate *E. coli* colonies, and for the examination of the antibiotic resistance profile. Containers were rinsed three times with water collected at each test point to minimise or totally eradicate any potential contaminants. In order to sample surface water, the sample bottle was carefully lowered horizontally into 30 cm depth in the river, with its mouth pointing upstream, while taking appropriate precautions to prevent floating or suspended debris. Finally, the samples were stored in a cooler box and transported to the Science and Technology Laboratories, Iraqi Ministry of Higher Education, for analysis within 24 h. This method prevents flocculation and microbial growth, which could have an impact on the outcome [75].

#### 3.2.1. Field and Laboratory of Physiochemical Measurements

Ranges of physical and chemical parameters were measured in the field, while others were conducted in the laboratory. Physical-chemical water quality parameters were analyzed using standard methods. This is described in another study submitted for publication.

#### 3.2.2. Faeces Sampling (Non-environmental)

Fecal samples were collected from individuals and animals as a potential source of contamination and to compare the antibiotic resistance patterns of *E. coli* with those of river samples, from the surrounding areas of the study sites and other random locations within Karbala city. Fifty fresh fecal samples were collected from human volunteers, animals (cats and birds), and from the ground of the livestock farms and poultry houses (cows and poultry). Every faecal sample was kept in phosphate-buffered saline solution with 15% glycerol added, and they were all placed in specialized plastic containers. In order to isolate *E. coli*, a recognized possible source of isolation, the samples were brought to the lab.

### 3.3. *E. coli* Isolation and Identification

*E. coli* was isolated and identified using standard microbiological methods. Samples were carefully mixed and serially diluted by adding 1 ml of sample to 9 ml of 0.9% DW. This was followed by another series of dilutions (10-1 to 10-6). The Membrane Filter Method (MF) was used for detection and quantification of *E. coli* using membrane filters (Sartorius, Germany) 47 mm in diameter and 45 μm pore size. The filters and instruments were sterilized prior to use for 15 min at 121°C. After filtration, the membrane filter was gently transferred and placed upside down, avoiding bubble formation onto *E. coli* chromogenic agar (CEA; Himedia, India) plates and incubated for 24 h at 37°C. The characteristic blue colonies that were diagnostic of *E. coli* colonies were confirmed by re-culturing on EMB agar (EMB; Himedia, India) for 24 h at 37°C. Colonies with metallic green sheen color were counted as *E. coli*. The re-streaking procedure was carried out repeatedly until pure cultures were produced. All samples were kept in Luria Bertani broth (LB; Himedia, India) with 20% glycerol (Carlo Erba, France) at -80°C until further examined. Morphology, Gram stain, and biochemical tests were used in this investigation to identify *E. coli*. Biochemical assays included (Indole; Methyl Red; Simmons’ Citrate; and Voges–Proskauer) tests [75], [76], [77], [78].

#### 3.3.1. DNA Extraction and PCR Gene Identification

Following the manufacturer’s instructions, the FavoPrep™ genomic DNA extraction Mini kit was used to create the template DNA for PCR (FAVOUR GEN biotech corp., Taiwan). A monoplex PCR targeting the *E. coli* housekeeping gene (*uidA*) with primers and amplification conditions [79], [80] verified the identity of presumed *E. coli* isolates. The target gene (Table 1) was amplified by PCR using an Mj Mini Gradient Thermal Cycler (Mupid-exu, Japan). The PCR was conducted in 25 μl of reaction mixture consisting of 5 μl DNA, 1X Green GoTaq buffer (pH 8.5), 0.5U of Taq DNA polymerase (Promega, USA), 1.65 mM of MgCl_2_, 220μM dNTP, and 0.24 μM of each primer. After being electrophoresed in a 1.5% agarose gel for 90 minutes at 90V, the amplified PCR products were visualized using Gel Documentation (Bio-Rad, USA).

**Table 1.**
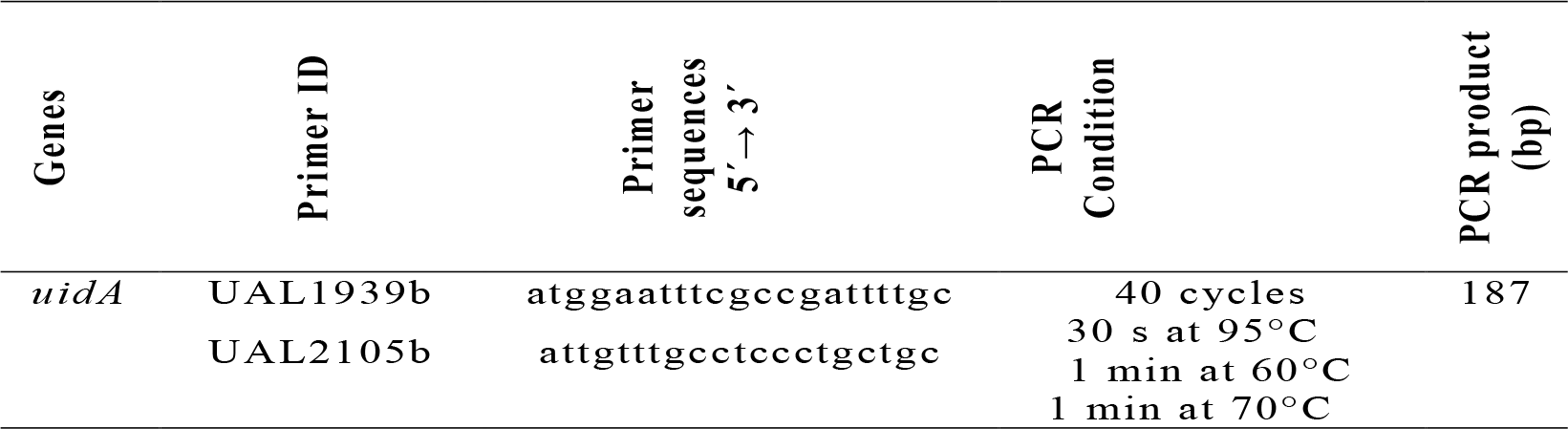
Primers used for PCR amplification and cycling conditions for selected gene.

### 3.4. Antibiotic Susceptibility Testing

The antibiotic resistance profiles of the *E. coli* isolates had been determined using the disc diffusion method, depending on recommendations of the Clinical and Laboratory Standards Institute [81]. A total of 12 typically used antibiotics (Himedia, India) from 11 classes were examined: penicillins [ampicillin (AMP) 10 μg], macrolide [azithromycin (AZM) 15 μg], cephalosporins [cefotaxime (CTX) 30 μg; ceftriaxone (CTR) 30 μg], amphenicol [chloramphenic (C) 30 μg], fluoroquinolones [ciprofloxacin (CIP) 5 μg], sulfonamides [Co-trimoxazole / sulpha /trimeth (COT) 25 μg], aminoglycoside [gentamicin (GEM) 10 μg], carbapenems [meropenem (MRP) 10 μg], quinolones [nalidixic acid (NA) 30 μg], tetracyclines [tetracycline (TE) 30 μg], glycylcycline [tigecycline (TGC) 15 μg]. The turbidity of fresh *E. coli* suspensions was adjusted to meet the McFarland standard of 0.5 ± 0.05 by UV-Vis Spectrophotometer (Sco-tech, Germany). A Mueller Hinton (MH; Himedia, India) agar plates was covered via cotton swabs to properly distribute the suspension. The plates were covered with antibiotic discs and incubated for the 24h at 37 ± 0.5°C. Each *E. coli* isolate’s zone of inhibition was examined using CLSI’s standards and interpretation criteria [81] as resistant (R), intermediate (I), and susceptible (S) [82], [83]. Multidrug resistance (MDR) was the designation given to an isolate if it demonstrated resistance to three or more distinct antibiotic classes [84].

#### 3.4.1. Multiple Antibiotic Resistance Index (MAR)

The following calculation was used to get the average MAR index of all *E. coli* isolates:

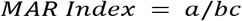

Where “a” is the aggregate antibiotic resistance score of all the *E. coli* isolates from one season or one site, “b” is the number of *E. coli* isolates tested, and “c” is total number of antibiotics that were tested on each isolate. When MAR index values ≥ 0.2 refer source of high risk of pollution where antibiotics are common used [85], [67].

### 3.5. Statistical Analysis

Microsoft Excel 2016 was used to enter raw data and create statistical summaries. Analysis of variance (ANOVA) was performed to determine the variance between antibiotics of river isolates and non-environmental isolates at a significance level of P < 0.05. Hierarchical cluster heat maps were used for both river isolates and non-environmental isolates, as well as for study sites and seasons. Spearman’s correlation analysis was used to find associations between water quality parameters and AREs. Previous analyses were performed using OriginPro 2024 software, USA.

## 4. Conclusions

This study aimed to assess the prevalence of AREs in the Euphrates River waters in different urban and agricultural areas in Iraq. Data show an increase in the contamination of river waters with AREs, which coincided with high rates of AREs in non-environmental samples. The results demonstrated that wastewater treatment plant discharges, agricultural activities, and livestock farming might lead to differences in the proportions of AREs. These data may be useful in developing specific regulations for AREs transmission and waste management in various urban and agricultural areas. This study provides baseline data for the continuous monitoring of natural water bodies in the future, in conjunction with improved livestock wastewater treatment and the development of sewerage networks. Through continuous monitoring, the overall impacts of animal wastewater treatment and sewerage network disinfection on AREs can be better understood. Furthermore, intervention strategies can be used to identify hotspots of multidrug-resistant bacteria, which will help reduce environmental contamination by AREs.

## Acknowledgments

The authors are thankful to the College of Science for Women, University of Baghdad, and the laboratories of the Ministry of Science and Technology.

## Conflicts of Interest

The authors declare no conflict of interest.

**Supplement**: Google My Map

